# Video Recording Can Conveniently Assay Mosquito Locomotor Activity

**DOI:** 10.1101/2020.02.28.970566

**Authors:** Maisa da Silva Araujo, Fang Guo, Michael Rosbash

**Author notes:** Laboratory of Entomology, Fiocruz Rondônia, Brazil and PGBIOEXP/PNPD, Federal University Foundation of Rondônia, Brazil. Department of Neurobiology, Key Laboratory of Medical Neurobiology of the Ministry of Health of China, Key Laboratory of Neurobiology, Zhejiang University School of Medicine, Hangzhou, Zhejiang 310058, China (Fang Guo).

## Abstract

*Anopheles gambiae* and *Aedes aegypti* are perhaps the best studied mosquito species and important carriers of human malaria and arbovirus, respectively. Mosquitoes have daily rhythms in behaviors and show a wide range of activity patterns. Although *Anopheles* is known to be principally nocturnal and *Aedes* principally diurnal, details of mosquito activity are not easily assayed in the laboratory. We recently described FlyBox, a simple tracking system for assaying *Drosophila* locomotor activity rhythms and thought that it might also be applicable to monitoring mosquito activity. Indeed, we show here that FlyBox can easily, conveniently, affordably and accurately measure the activity of *Anopheles* as well as *Aedes* over several days. The resulting profiles under light-dark as well as constant darkness conditions are compatible with results in the literature, indicating that this or similar systems will be useful in the future for more detailed studies on a range of insect species and under more diverse laboratory conditions.

## Introduction

The female mosquito is the principal vector of several vector-borne diseases affecting human and other animals in tropical and temperate parts of the globe ^1^. Most adult female mosquitoes are haematophagous, namely, they need to take a blood-meal for maturation of female oocystes ^2^. This blood feeding allows the transmission of several zoonotic and human disease agents ^3^, such as parasites (malaria and filariasis) and arboviruses (yellow fever, dengue, chikungunya and Zika virus).

Among the three subfamilies of Culicidae, two are of medical interest: the Anopheline, with the most important genus, *Anopheles;* and the Culicinae, with principally the genera *Aedes*, and *Culex* ^4^. *Anopheles gambiae* and *Aedes aegypti* are perhaps the best known species and are carriers of human malaria and arbovirus, respectively. *Anopheles gambiae* is the primary vector for African malaria, which is caused by parasites from the *Plasmodium* genus ^5^. There were an estimated 435,000 malaria deaths worldwide in 2017, and most were in Africa ^6^. *Aedes aegypti* is a global vector for many human diseases, such as yellow fever, dengue, chikungunya and Zika ^7^. Over half the world population is at the risk of dengue and chikungunya infections ^8^, and that of other arboviruses keeps on increasing globally.

Mosquitoes have daily rhythms that restrict their activity, such as flight, mating, sugar or blood-meal feeding and oviposition, to specific hours of the day. The cause of these daily rhythms is an endogenous circadian clock; it can be synchronized to external cues, such as light, temperature and food ^9^. The relationship between the circadian clock and the different mosquito behaviors is particularly important to their vector capacity. For example, host-seeking is crucial to vector efficiency and is influenced by the circadian clock. Importantly, different mosquito species show a wide range of activity patterns, including diurnal, crepuscular, and nocturnal ^10^.

Not surprisingly, disease transmission follows the activity patterns of the specific mosquito vectors. Knowledge of mosquito behavior is therefore important for understanding the dynamics of mosquito-host contact, i.e., mosquito behavior influences the intimacy of contact with humans ^11^. Developments in automated tracking techniques allow recording and observation of some important mosquito behaviors, and software programs can quantify them. These techniques therefore produce data that can aid in the interpretation of mosquito behavioral details^12,13^.

Laboratory measurements of mosquito locomotor patterns was initiated with acoustic measurements in sound-proof flight cages ^14,15,16^ and has recently been done with break-beam technology^17,18,19^. To monitor more precisely the movement and sleep of adults *Drosophila* flies, Guo *et al*.^20^ used an automated video recording assay instead of the beam-break *Drosophila* activity monitor (DAM) system. Guo *et al*.^21^ then developed a simple modification, called FlyBox, which can video record real time fly behavior in a 96-well plate. The FlyBox system is an independent light-tight box equipped with entrainment lights and video recording to detect activity/sleep patterns. For the *Drosophila* experiments, there were also optogenetic LEDs in FlyBox, which allows the activation or inhibition of neurons noninvasively and detects the corresponding behavioral response. The simpler version of FlyBox used in this paper did not contain the optogenetic LEDs.

Considering that this new system is easy, convenient, affordable and produces reliable data, we used FlyBox to assess the locomotor activity patterns of two important mosquito tropical disease vectors, *An. gambiae* and *Ae. aegypti*. In addition, male and female locomotor activity patterns were compared to assess differences between the sexes. These differences might be exploited for the development and/or evaluation of vector control strategies. The validation of this relatively new system to study locomotor activity beyond *Drosophila* may be useful for many different insect species.

## Results

We monitored the locomotor activity patterns of *An. gambiae* and *Ae. aegypti* using FlyBox (Fig. 1 and see Supplementary Video), a relatively new behavior monitoring system first developed for *Drosophila* ^21^. We evaluated the ability of the FlyBox basic version (without optogenetics LEDs; see below) to assay mosquito activity by comparing two important tropical disease vectors, *An. gambiae* and *Ae. aegypti*. The mean locomotor activity of *An. gambiae* and *Ae. aegypti* females and males was first monitored for four full days in 12 h light:12 h dark (LD) conditions (Fig. 2A).

**Figure 1.**
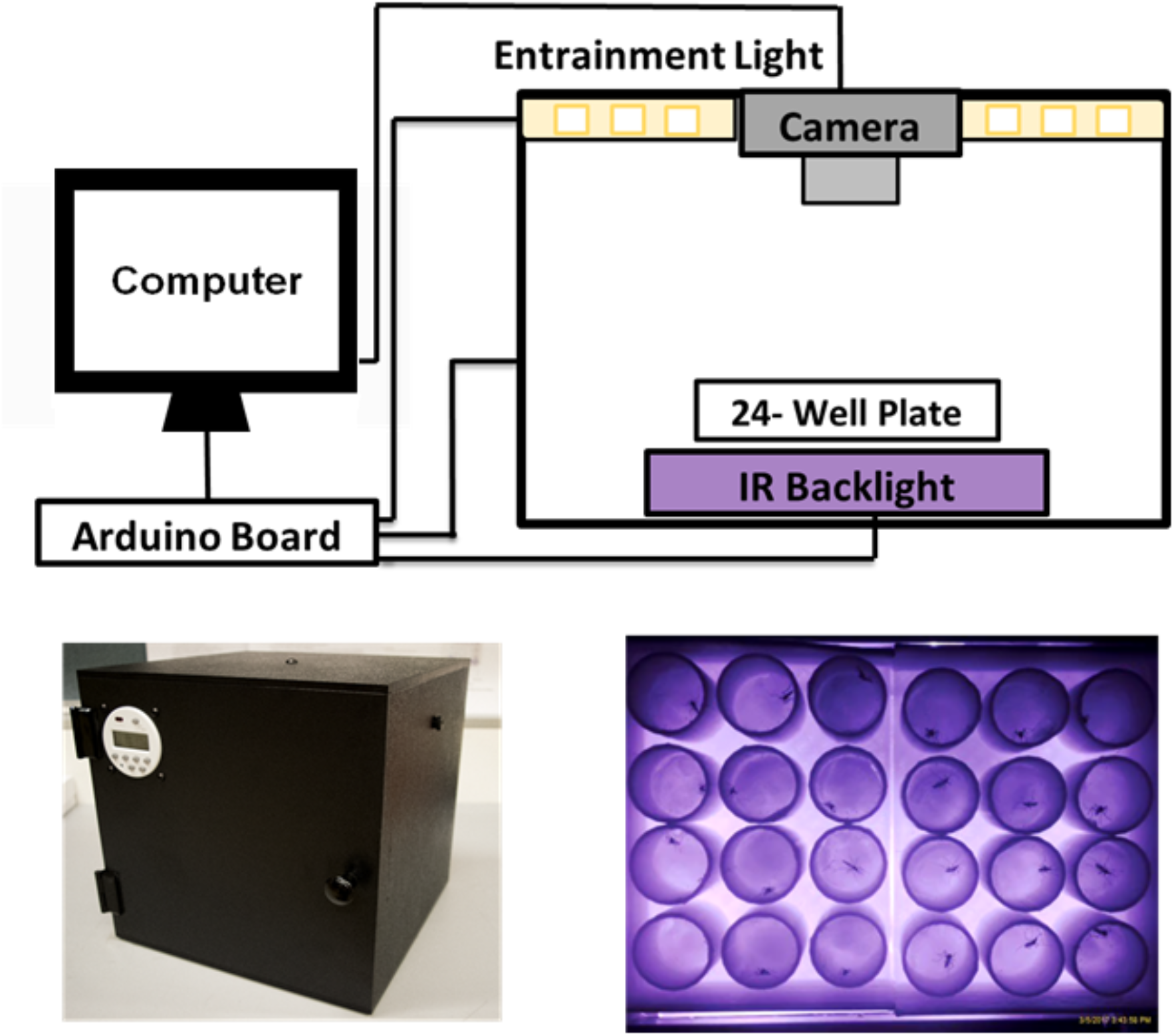
Schematic of FlyBox. Illustration of FlyBox schematic (upper panel) and the front view of FlyBox (lower left panel) were shown. The example of recording image used for mosquito behavior monitoring is provided in lower right panel.

**Figure 2.**
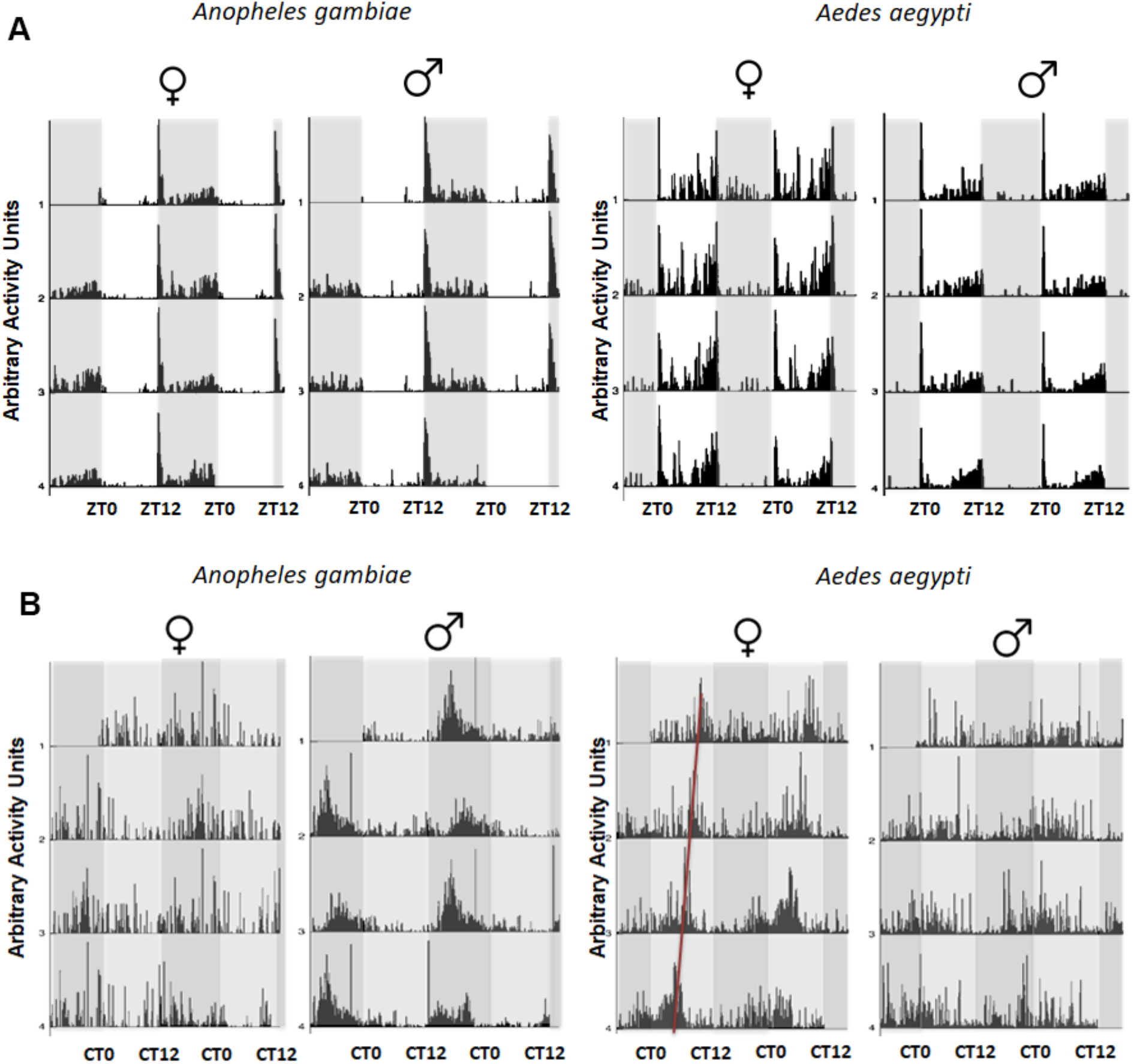
(A) Locomotor activity of *Anopheles gambiae* and *Aedes aegypti* mosquitoes under LD conditions. Double-plotted actograms show the average locomotor activity of individually monitored males and females of both species under cycles of 12 h light and 12 h of dark (LD) over four days. ZT indicates zeitgeber time. Gray shading indicates darkness. (B) Locomotor activity of *Anopheles gambiae* and *Aedes aegypti* mosquitoes under DD conditions. Double-plotted actograms show the average locomotor activity of individually monitored males and females of both species under constant dark (DD) over four days. Light gray background represents “subjective day” and dark gray background “subjective night”. n= 11-12.

*Anopheles gambiae* activity is largely restricted to the nighttime as expected for this nocturnal vector (p<0.05 for the *Anopheles gambiae* female group and p<0.01 for the *Anopheles gambiae* male group by student’s T-test in Fig. 3), whereas the diurnal vector *Ae. aegypti* is active mainly during the day (p<0.05 for the *Ae. aegypti* female group and p<0.01 for *Ae. aegypti* male group by student’s T-test in Fig. 3). The rhythms of both species are therefore quite similar except for reversing day and night (Fig. 3). The nocturnal rhythms of *An. gambiae* show a pronounced activity peak at dusk, whereas the daytime activity of *Ae. ae*gypti shows pronounced peaks at dawn as well as dusk (Fig. 2A).

**Figure 3.**
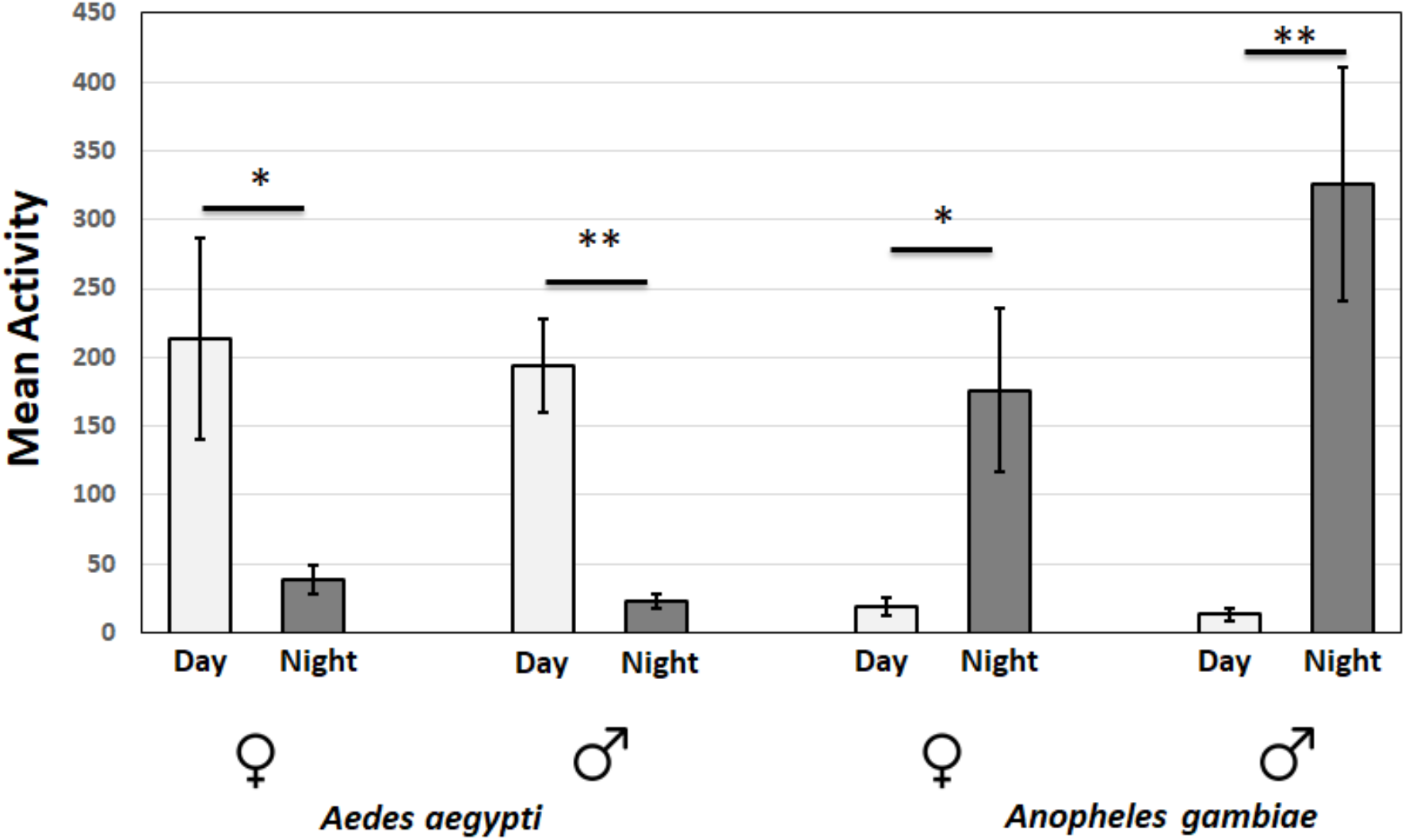
Comparisons of daytime and nighttime mean activity of *Anopheles gambiae* and *Aedes aegypti* mosquitoes under LD conditions. Light gray bars represent “day” and dark gray bars represent “night”. n= 10-12. * p<0.05 and ** p<0.01 by student’s T-test.

Activity monitoring was then continued in constant darkness (DD; Fig. 2B). This is the first time that the activity of these two species has been compared under both LD and DD conditions in a single study. The pronounced *Ae. aegypti* dawn activity peak disappeared even in the first day of DD (Fig. 2B). On the other hand, there is an evening peak of activity in both species that continues in DD with a period about 22h. (See the red line in the Fig. 2B female plot). There are no notable differences in circadian period between sex and species (Figs. 2B and 4A and B). Rhythmicity was observed in 95% of females and about 90% males of *An. gambiae*, whereas 75% females and 81% males of *Ae. aegypti* were rhythmic (Fig. 4C and D).

**Figure 4.**
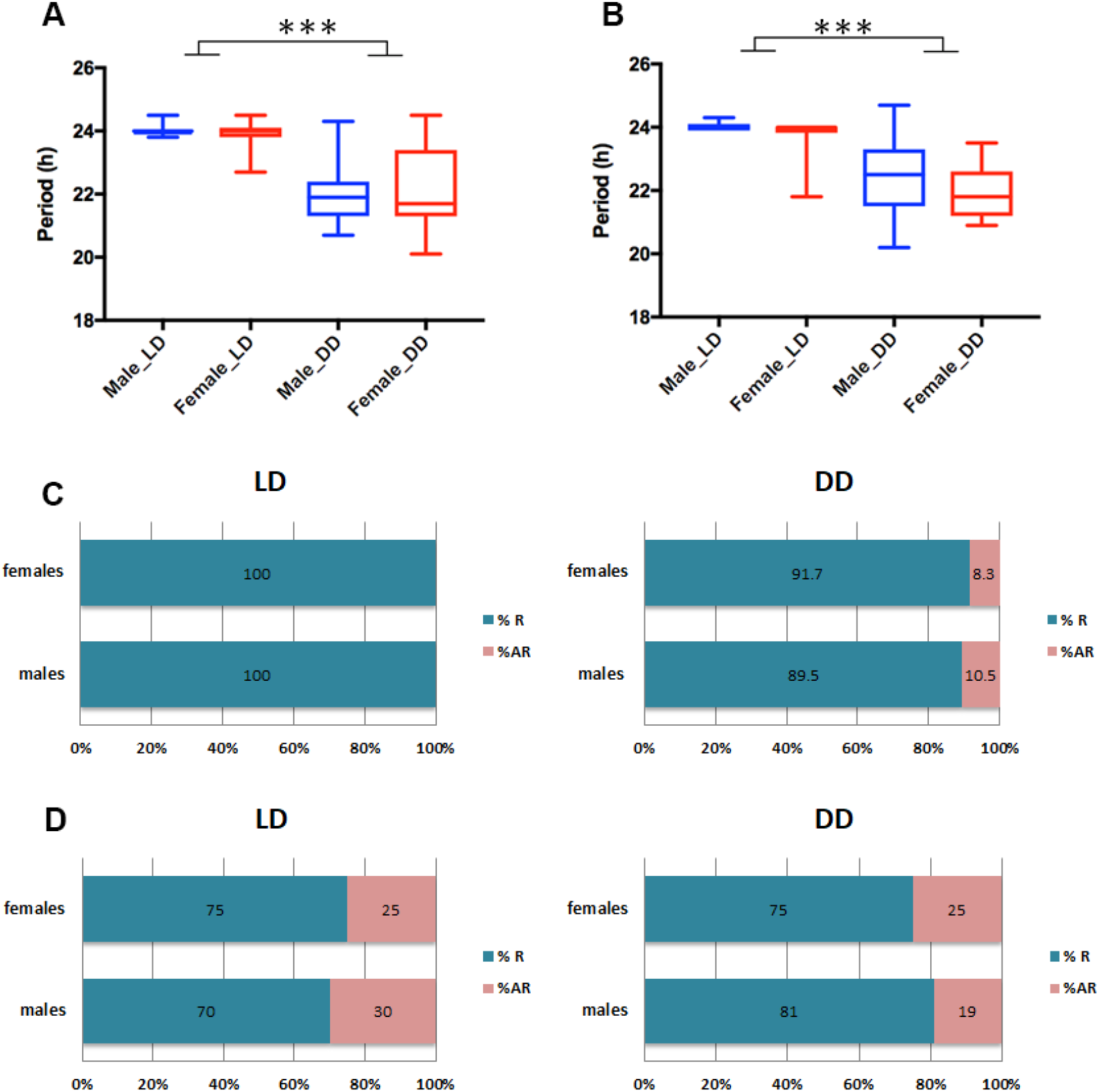
Analysis of circadian period and quantification of rhythmic and arrhythmic male and female *Anopheles gambiae* and *Aedes aegypti*. There are no apparent sex differences, and LD periods are clamped at 24hr as expected. (A) Mean ± SEM of *Anopheles gambiae* and (B) *Aedes aegypti* period in hours. *** p<0.001, Ordinary one-way ANOVA. n = 18-23. *Anopheles gambiae* (C) and *Aedes aegypti* (D) mosquitoes under LD and DD conditions. % R (blue) represents the percentage of rhythmic flies. % AR (pink) represents the percentage of arrhythmic mosquitoes. n = 23-24.

The somewhat short period of about 22h has also been observed by others ^14,18^. In our hands however, the period length of males was not different from females (p > 0.05 by using student’s T-test in Fig. 4A and B), in contrast to a previous publication ^18^. These authors reported that there are sex-specific differences in the circadian period length of *An. gambiae*, namely, the locomotor activity of males is earlier than that of females under entrained conditions ^18^. This is because the activity phase is a direct measure of the period length of the underlying endogenous circadian clock ^22^.

To highlight differences between species, the LD behavior patterns were compared between females of *An. gambiae* and females of *Ae. aegypti* (Fig. 5A). As suggested by examining the daily plots (Fig. 2), *An. gambiae* has a nearly unimodal pattern: there is a sharp, major peak of activity directly after lights-off, which rapidly declines. There is also some modest subsequent activity, which progressively increases during the second half of the night, between ZT19 and ZT1. In contrast, *Ae. aegypti* has a clear bimodal activity pattern, with morning and evening activity peaks. The morning peak is certainly a startle response to lights on, as there is no indication of anticipatory activity and this peak is absent in DD (Fig. 2B). The evening peak is centered on the light-to-dark transition and in contrast to the morning peak shows a clear anticipation of that transition.

**Figure 5.**
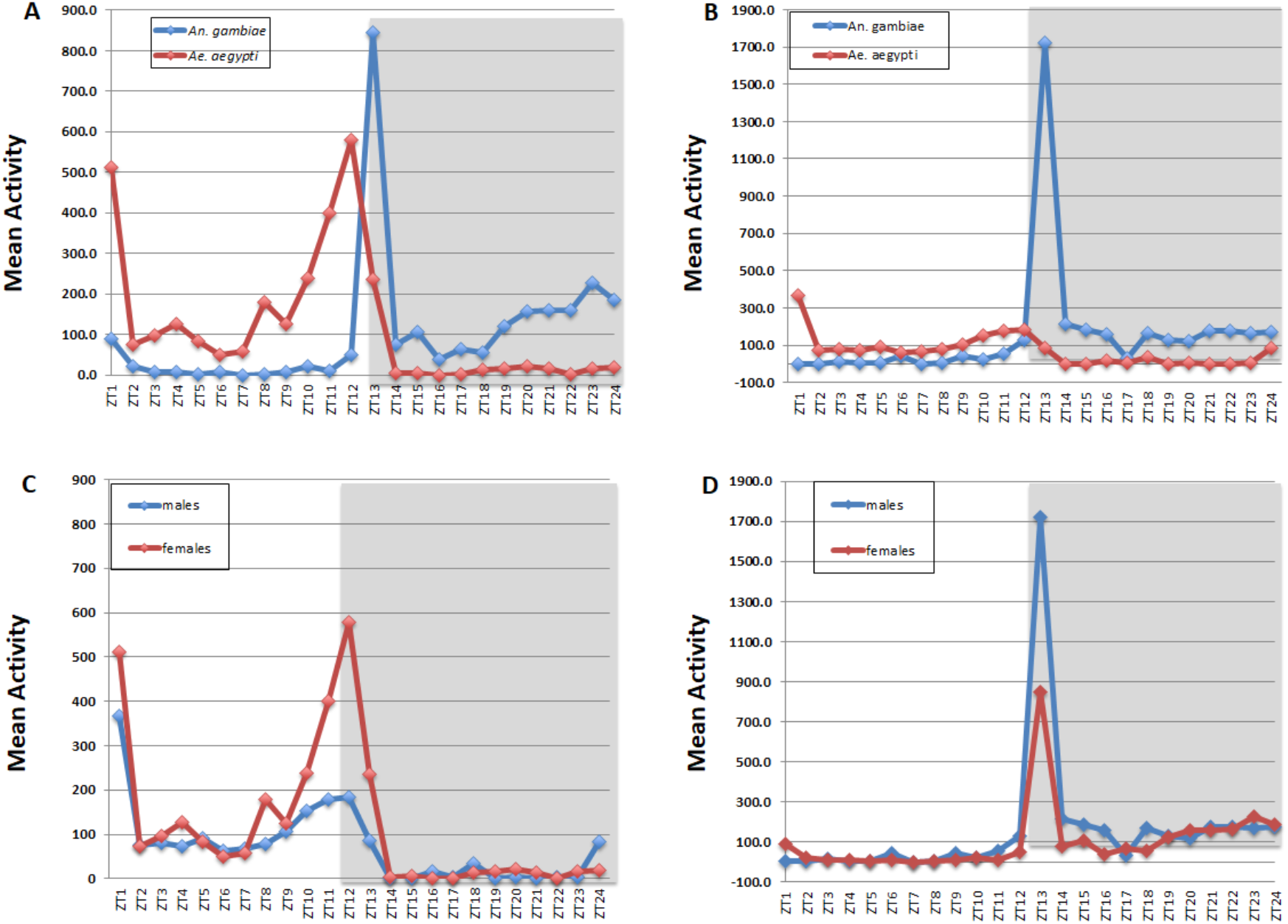
Mean locomotor activity profiles. (A) Profiles of females, *Anopheles gambiae* vs *Aedes aegypti;* (B) Profiles of males, *Anopheles gambiae* vs *Aedes aegypti;* (C) Profiles of *Aedes aegypti*, males vs females; (D) Profiles of *Anopheles gambiae*, males vs females. The graphs show the mean activity of 23-24 mosquitoes during each ZT hour over 4 days in LD12:12.

The same species comparison of males is rather similar to that of females, except that the overall activity of *Ae. aeg*ypti males is low compared to *An. gambiae* males (Fig. 5B). Nighttime activity of *Ae. aegypti* males is very low, and *An. gambiae* males do not respond to lights-on (Fig. 5B). The low activity of *Ae. aegypti* males is also apparent compared to *Ae. aegypti* females (Fig. 5C).

Lower male activity is also the major difference between *Ae. aegypti* males and females. As described above, both sexes have prominent activity at dawn and a prominent evening activity peak in anticipation of lights off. Notably, the dawn activity is absent in DD (Fig. 2B), consistent with the interpretation that it is a startle response to lights-on. Male and female *An. gambiae* also have very similar activity patterns with a very modest activity increase in anticipation of the lights-off transition (Fig. 5D).

Lastly, we assayed the activity of a newly established *Anopheles* colony under LD conditions and without blood feeding. Males and females were separated following emergence, and the locomotor activity of individual virgin mosquitoes was then recorded for a week with FlyBox. The mean activity of virgin males and females 6 days after emergence was very similar (p=0.7258 by student’s T-test in Fig. 6). However, virgins revealed significantly more activity with a pronounced activity peak at dusk when compared with inseminated mosquitoes (p<0.001 by student’s T-test for virgin group in Fig. 6 vs inseminated group in Fig. 5D). This same marked activity increase was observed by Jones and Gubbins ^23^ using the acoustic autograph technique in *An. gambiae* (Pala strain females). The dusk activity peak of virgins suggests swarming and mating behavior and also corresponds to the time of host-seeking behavior ^23^. This activity peak is greatly reduced later in the night (Fig. 5D), consistent with the known timing of these two functions as well as oviposition. The virgins also have an additional activity peak at dawn.

**Figure 6.**
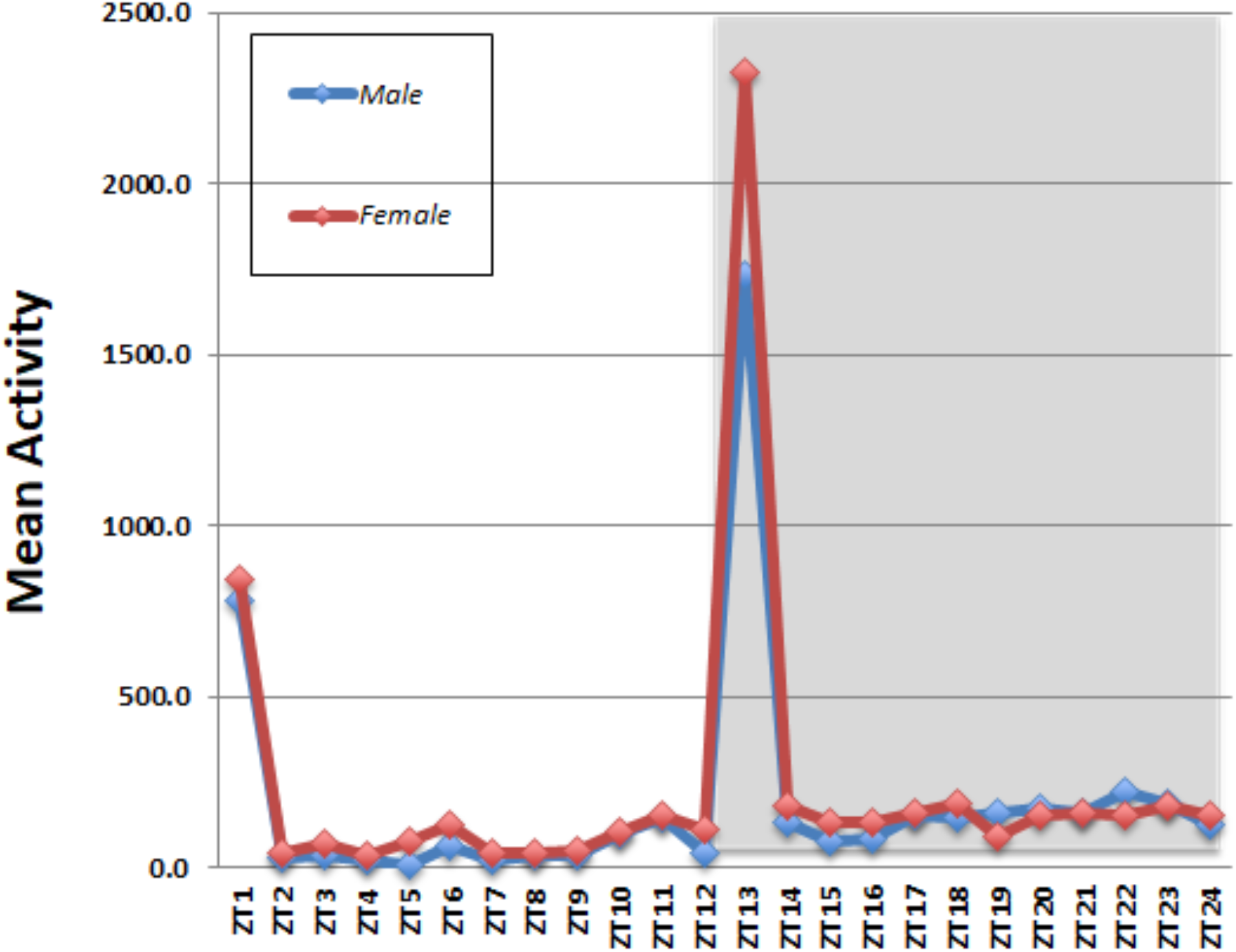
Mean locomotor activity profiles of virgin *Anopheles*, males vs females. The graphs show the mean activity of 23-24 mosquitoes during each ZT hour over 4 days in LD12:12.

## Discussion

In this study, we used the FlyBox to assay mosquito activity. The 24-well plate system allowed us to automatically and simultaneously analyze 24 individual mosquitoes for a full six days under both LD and DD conditions. Although new software for tracking multiple individuals in a larger arena is becoming available, behavioral observations on isolated individuals is still standard practice and more straightforward. However, this single mosquito strategy would have been much more laborious and time-consuming without FlyBox. Mosquito locomotor activity had been previously monitored in the laboratory by acoustic measurements, for example in sound-proof flight cages ^15,16,24–27^ or by utilizing beam-break technology ^18,19,28–32^. The FlyBox system was adapted from insights gained from previous approaches to the study of *Drosophila* locomotor activity behavior ^21^.

The advantages of FlyBox compared to the traditional DAM system (DAM) are i) spatial and movement resolution is higher. The DAM system uses a single infrared beam break system. Consequently, it loses spatial and temporal resolution due to its intrinsically simpler design; ii) shorter preparation time. It is easier to load food in the 24-well plates than in the DAM glass behavior tubes. The disposable plates also do not need to be cleaned or autoclaved like the tubes; iii) FlyBox is compact (11 inch cube) and has its own entrainment lights, which avoids the requirement of incubators and their separate entrainment light cycles. As a consequence, FlyBox is easily transported and can be used in a temperature and humidity-controlled room such as in an insectary or even just on a benchtop; iv) it is cost effective and efficient and can be easily built. For example, the automatic analysis of behavior enables the use of an inexpensive ($70-80) webcam (1280 x 720 pixels; 1920 x 1080 pixels for video).

The drawbacks are: i) 24-well plates detect fewer mosquitoes than the standard 32 glass behavior tubes of the DAM system; ii) The limited space within each well of the 24-well plate is non-physiological and almost certainly restricts the natural (flight) activity of the insect (Fig. 1 and see Supplementary Video) albeit less severely than the DAM tubes. Indeed, video (see Supplementary Video) shows that the mosquitoes are jumping and attempting flying. However, this drawback underscores the extent to which the circadian clock must regulate the temporal patterns so that they still resemble previous data in the literature.

*Anopheles gambiae* are primarily night biters ^33,34^. Our data indeed show that *An. gambiae* mosquitoes are predominately nocturnal, and the evening peak is consistent with flight activity for mating. This broad activity peak corresponds to the well-known mating swarm behavior ^14,35,36^. We attribute the pronounced male activity peak to the need to form swarms; they are composed almost entirely of males, and single females briefly fly and leave the swarm in copula. Copulation is generally fast, less than 20 sec.^36^, which means that flight activity might be less for females than for males. Nocturnal activity is also consistent with female host-seeking and with inseminated female oviposition ^37,38^. Experiments that recorded biting activity and oviposition of *An. gambiae* showed that both activities are essentially nocturnal; oviposition peaks just after sunset, whereas biting activity is maximal after midnight and in the hours before dawn ^38^. A similar differential behavioral pattern between male and female mosquitoes was documented for *An. gambiae* Pimperena S form and for Mali-NIH M form colonies using the DAM system^18^.

Because most of our experiments were done with inseminated females without blood feeding, it is likely that the pronounced activity peak shortly after lights off essentially corresponds to host-seeking. The second more gradual rise in activity would then correspond to biting activity and sugar feeding as described in an analysis of sugar feeding frequency in males and females of *An. gambiae* ^39^. These insects are also very inactive in the light ^18,40^.

*Aedes aegypti* have already been reported as a diurnal vector with bimodal rhythms under laboratory conditions ^41,42–44^. Trimodal patterns have also been reported in an automatic recording device ^28^. The morning activity peak was common in both sexes but disappears under free-running (DD) conditions (Fig. 2), indicating a response to light as previously observed ^19,43,44^ In contrast, the major evening peak in *Ae. aegypti* females continues under DD conditions with males showing reduced activity at this same time of day. This morning and late afternoon activity are consistent with reports that this is when *Aedes* mates ^45^. However, males of this genus do not require swarms like *Anopheles* but are generally able to mate in small groups or in single pairs ^46^. The activity peaks at sunrise and during the day are consistent with the flight activity of host-seeking and sugar feeding-cycles ^44^. Moreover, the lights-off peak in *Ae. aegypti* shows a characteristic nocturnal anticipation, similar to the diurnal insect *Drosophila melanogaster* ^47^. This may only be a feature of diurnal insects as nocturnal anticipation is not clear in *An. gambiae*.

To summarize, we used FlyBox to compare different circadian activity patterns between *An. gambiae* and *Ae. aegypti*. The results show that automatic recording of mosquito activity is possible in this small, simple and inexpensive monitoring system; it costs approximately $500 for the parts. We further suggest that FlyBox will be able to increase the number of laboratory studies addressing different insect vectors of human diseases and may even facilitate drug screening to combat these vectors. For example, the single animal FlyBox wells should facilitate drug screening at different sub-lethal concentrations. Moreover, the portability of FlyBox should make it possible to assay mosquito activity in remote locations. This should more easily allow the study of behavioral shifts in wild populations, where mosquitoes have been under selective pressure due to agents like insecticides.

## Methods

### Mosquito rearing

Mosquitoes used in this study were from a laboratory colony of *An. gambiae* sensu stricto strain G3 and *Ae. aegypti* Rockefeller strain maintained at Laboratory of Dr. Flaminia Catteruccia from Harvard T. H. Chan School of Public Health, Department of Immunology and Infectious Diseases. The colonies were reared under standard conditions (26°C ± 1°C, 70% ± 1% humidity and 12 h: 12 h LD photoperiod, without dawn and dusk transitions). For colony maintenance, 5-7-day-old females were provided a blood meal of human blood weekly (Research Blood Components, Boston, MA) using an artificial membrane feeding system (Hemotek). For all experiments, mosquitoes were pre-synchronized from pupae to adult stage under the same conditions of temperature, humidity and photoperiod described above in a Precision Scientific Incubator Mod 818 the Rosbash laboratory. Female and male adults were maintained into same cage for insemination for 4-5 days and they were supplied with a basic diet of 10% sucrose solution on cotton wicks according to standard rearing conditions ^48^. All mosquitoes used for experiments were 4-7 days of age, mated (no virgins), and females were not blood feeding.

### Behavioral analysis

Circadian locomotor activity rhythms of *An. gambiae* and *Ae. aegypti* were recorded automatically using a new version of the automated behavior recording system FlyBox. It was previously optimized for *Drosophila* circadian rhythms and sleep as briefly described ^21^. In somewhat more detail, FlyBox is an 11 inch cube box made from black Acrylonitrile Butadiene Styrene (ABS). It is fully light-tight to prevent any light contamination during constant darkness experiments. In the experiments shown here, FlyBox was kept inside a climate-controlled insectary under the conditions described, but it can be used anywhere as long as it is connected to a computer to control its parameters and record data. FlyBox contains three main features: an entrainment light (5000K daylight white LED to fully mimic daylight), an infrared camera to record locomotor activity and a plate platform (Fig. 1). The FlyBox can also be configured to include optional red and green LEDs for optogenetics, which were not used in this study. A 24-well plate sufficiently large for mosquitoes replaced the 96-well plate used for fruit flies ^21^. The plate platform, made of white acrylic transparent to infrared light, is positioned in the center of the box, and the entrainment light is placed in the back to provide even illumination. A pair of 850nm infrared LEDs are placed underneath the plate platform to enable continuous observation by the camera through day and night. The entrainment light is controlled by a BuckPuck LED driver connected to an Arduino Uno, while the intensity of the infrared LEDs are controlled by a BuckPuck connected to a 5k pot. The FlyBox camera is also directly connected to the computer for data capture. Please email the authors for more detail about parts and construction.

There are three basic steps to monitoring mosquito circadian locomotor activity in FlyBox.

1. Preparation of the 24-well plate: Adult male and female mosquitoes were removed from cages with an automatic insect aspirator and anesthetized on ice until individually placed into wells using forceps. At the bottom of each well is a piece of cotton soaked in 10% sucrose solution. This allowed the mosquitoes to eat *ad libitum* during the observation periods. After covering the 24-well plate with a transparent plastic film and poking five holes over each well to promote air circulation, it was placed on the platform inside the FlyBox.
2. Configuring and running FlyBox. The fully loaded 24-well plate can be seen in real time with the FlyBox camera, which is connected to a computer (Fig. 1). Locomotor activity starts recording when the software (WebCamImageSave) is opened, and an image is captured every 10 seconds. All recordings were in a 12h:12h LD cycle or in DD conditions. Time of day in LD is reported in 24hr Zeitgeber Time (ZT); ZT12 is defined as the time of lights off under the LD cycle, and ZT0 defined as lights-on (dawn). Time of day in DD is in traditional circadian time (CT). Two replicates of the behavior recording were done with 12 males and 12 females of each species. Mosquitoes were monitored for at least 7 days, of which the first two days were just to synchronize under LD conditions. These two days and the last day of consistent activity were excluded from the analysis. Excluding the first two days should mitigate against any effects of the anesthesia from the ice-cooling. Two independent behavioral experiments were conducted for each species: one under LD 12:12 for seven days (LD experiment) and the other under LD 12:12 for two days followed by constant darkness for five days (DD experiment). The data of individuals that died during the experiments were excluded, and the analysis was carried out comparing the activity data of males vs. females and *An. gambiae* vs. *Ae. aegypti* mosquitoes.
3. FlyBox data processing and analysis. This is almost identical to analysis from the traditional DAM system ^21^. The pictures from the WebCamImageSave software were saved every 10 seconds and then used by PySolo to calculate the travelled distance of each mosquito. The data then were converted into a .txt file to run a MATLAB (MathWorks) program called Sleep and Circadian Analysis MATLAB Program (SCAMP) from the Griffith lab ^49^. Activity is defined by >5 pixel change of mosquito positions between pictures captured every 10 seconds. The total distance travelled in 1 min was calculated by MATLAB for plotting. Rhythmicity Index (RI) derived from the autocorrelation function was used to calculate DD rhythmicity, and mosquitoes with a RI lower than 0.15 were considered arrhythmic ^50^.

## Acknowledgements

We thank A. Yu for help with the building and the description of FlyBox. Dr. Maisa Araujo’s work was funded by PDE/CNPq.

## Author contributions statement

MA, FG and MR conceived and designed the experiments. MA collected the data and performed the analysis. FG contributed analysis tools and designed Figure 1. MR, FG and MA wrote the paper.

## Data availability statement

The datasets generated during and/or analyzed during the current study are available from the corresponding author on reasonable request.

## References

1 Bigoga, J. D., Manga, L., Titanji, V. P., Coetzee, M. & Leke, R. G. Malaria vectors and transmission dynamics in coastal south-western Cameroon. Malar. J. 6, 5–33, doi:10.1186/1475-2875-6-5 (2007).

2 Pina, I. G. & Fonseca, A. H. Comportamento de *Aedes aegypti* L., (Diptera: Culicidade) alimentados artificialmente com sangue de diferentes espécies de doadores. Rev. Patol. Trop. 28, 64–71 (1999).

3 Coker, H. A. B., Chukwuani, C. M., Ifudu, N. D. & Aina, B. A. The malaria scourge concepts in disease management. Nig. J. Pharm. 32, 19–46 (2001).

4 Berenger, J. M. & Parola, P. Arthropod Vectors of Medical Importance in Infectious Diseases (Fourth Edition) 104–112 (Cohen, J. et al. 2017).

5 White, G. B. *Anopheles gambiae* complex and disease transmission in Africa. Trans. R. Soc. Trop. Med. Hyg. 68, 278–301, doi:10.1016/0035-9203(74)90035-2 (1974).

6 World Health Organization. World Malaria Report 2018 https://www.who.int/malaria/publications/world-malaria-report-2018/report/en/ (World Health Organization, Geneva, 2018).

7 Powell, J. R. Perspective Piece Mosquito-Borne Human Viral Diseases: Why *Aedes aegypti*? Am. J. Trop. Med. Hyg. 98, 1563–1565 (2018).

8 Kraemer, M. U. et al. The global distribution of the arbovirus vectors *Aedes aegypti* and *Ae. albopictus*. Elife 4, e08347, doi:10.7554/eLife.08347 (2015).

9 Hardin, P. E. Molecular genetic analysis of circadian timekeeping in *Drosophila*. Adv. Genet. 74, 141–173, doi:10.1016/B978-0-12-387690-4.00005-2 (2011).

10 Clements, A. The Biology of Mosquitoes. Vol. 2 (1999).

11 Wernsdorfer, W. H. & McGregor, I. Malaria: Principles and Practice of Malariology. illustrated edn, 1818 (Churchill Livingstone, 1988).

12 Spitzen, J. & Takken, W. Keeping track of mosquitoes: a review of tools to track, record and analyse mosquito flight. Parasit. Vectors 11, 123–134 (2018).

13 Wilkinson, D. A., Lebon, C., Wood, T., Rosser, G. & Gouagn, L. C. Straightforward multi-object video tracking for quantification of mosquito flight activity. J. Insect. Physiol. 71, 114–121 (2014).

14 Jones, M. D. R. & Gubbins, S. J. Changes in the circadian flight activity of the mosquito *Anopheles gambiae* in relation to insemination, feeding and oviposition. Physiol. Entomol. 3, 213–220, doi.org/10.1111/j.1365-3032.1978.tb00151.x (1978).

15 Nayar, J. K. & Sauerman, D. M., Jr. The effect of light regimes on the circadian rhythm of flight activity in the mosquito *Aedes taeniorhynchus*. J. Exp. Biol. 54, 745–756 (1971).

16 Jones, M. D. R. The automatic recording of mosquito activity. J. Ins. Physiol. 10, 343–351 (1964).

17 Sampaio, V. S. et al. What does not kill it makes it weaker: effects of sub-lethal concentrations of ivermectin on the locomotor activity of *Anopheles aquasalis*. Parasit. Vectors 10, 623–633 (2017).

18 Rund, S. S., Lee, S. J., Bush, B. R. & Duffield, G. E. Strain- and sex-specific differences in daily flight activity and the circadian clock of *Anopheles gambiae* mosquitoes. J. Insect. Physiol. 58, 1609–1619, doi:10.1016/j.jinsphys.2012.09.016 (2012).

19 Gentile, C., Rivas, G. B., Meireles-Filho, A. C., Lima, J. B. & Peixoto, A. A. Circadian expression of clock genes in two mosquito disease vectors: cry2 is different. J. Biol. Rhythms 24, 444–451, doi:10.1177/0748730409349169 (2009).

20 Guo, F. et al. Circadian neuron feedback controls the *Drosophila* sleep–activity profile. Nature 536, 292–297 (2016).

21 Guo, F., Chen, X. & Rosbash, M. Temporal calcium profiling of specific circadian neurons in freely moving flies. Proc. Natl. Acad. Sci. USA 114, E8780–E8787, doi:10.1073/pnas.1706608114 (2017).

22 Dunlap, J. C., Loros, J. J., Decoursey, P. J. Chronobiology: Biological Timekeeping. Sinauer Associates, Sunderland, MA. 2004.

23 Jones, M. D. R. & Gubbins, S. J. Modification of circadian flight activity in the mosquito *Anopheles gambiae* after insemination. Nature 268, 731–732 (1967).

24 Rowley, W. A., Jones, M. D., Jacobson, D. W. & Clarke, J. L. A microcomputer-monitored mosquito flight activity system. Ann. Entomol. Soc. Am. 80, 534–538 (1987).

25 Bennet-Clark, H. C. A particle velocity microphone for the song of smal insects and other acoustic measurements. J. Exp. Biol. 108, 459–463 (1984).

26 Peterson, E. L. The temporal pattern of mosquito flight activity. Behaviour 72, 1–25 (1979).

27 Jones, M. D. R., Cubbin, C. M. & Marsh, D. The circadian rhythm of flight-activity of the mosquito *Anopheles gambiae:* The light-response rhythm. J. Exp. Biol. 57, 337–346 (1972).

28 Kawada, H. & Takagi, M. Photoelectric sensing device for recording mosquito host-seeking behavior in the laboratory. J. Med. Entomol. 41, 873–881, doi:10.1603/0022-2585-41.5.873 (2004).

29 Chiba, Y., Uki, M., Kawasaki, Y., Matsumoto, A. & Tomioka, K. Entrainability of circadian activity of the mosquito *Culex pipiens pallens* to 24-hr temperature cycles, with special reference to involvement of multiple oscillators. J. Biol. Rhythms 8, 211–220, doi:10.1177/074873049300800304 (1993).

30 Shinkawa, Y. et al. Variability in circadian activity patterns within the *Culex pipiens* complex (Diptera: Culicidae). J. Med. Entomol. 31, 49–56, doi:10.1093/jmedent/31.1.49 (1994).

31 Chiba, Y. & Tomioka, K. Entrainability of diphasic circadian activity of the mosquito, *Culex pipiens molests*, to 24-hour light-dark cycles: A physiological significance of critical light-dark ratio. Zool. Sci. 8, 211–220 (1992).

32 Kasai, M. & Chiba, Y. Effects of optic lobe ablation on circadian activity in the mosquito, *Culex pipiens pallens*. Physiol. Entomol. 12, 59–65 (1987).

33 Rund, S. S., Gentile, J. E. & Duffield, G. E. Extensive circadian and light regulation of the transcriptome in the malaria mosquito *Anopheles gambiae*. BMC Genomics 14, 218–237, doi: 10.1186/1471-2164-14-218 (2013).

34 Bockarie, M. J. et al. The late biting habit of parous *Anopheles* mosquitoes and pre-bedtime exposure of humans to infective female mosquitoes. Trans. R. Soc. Trop. Med. Hyg. 90, 23–25, doi:10.1016/s0035-9203(96)90465-4 (1996).

35 Manoukis, N. C. et al. Structure and dynamics of male swarms of *Anopheles gambiae*. J. Med. Entomol. 46, 227–235, doi:10.1603/033.046.0207 (2009).

36 Charlwood, J. D. & Jones, M. D. R. Mating behaviour in the mosquito, *Anopheles gambiae* s.l. I. Close range and contac behaviour. Physiol. Entomol. 4, 111–120, doi:10.1111/j.1365-3032.1979.tb00185.x (1979).

37 Sumba, L. A. et al. Daily oviposition patterns of the African malaria mosquito *Anopheles gambiae* Giles (Diptera: Culicidae) on different types of aqueous substrates. J. Circadian Rhythms 2, 6, doi:10.1186/1740-3391-2-6 (2004).

38 Haddow, A. J. & Ssenkubuge, Y. Laboratory observations on the oviposition-cycle in the mosquito *Anopheles (Cellia) gambiae* Giles. Ann. Trop. Med. Parasitol. 56, 352–355, doi:10.1080/00034983.1962.11686130 (1962).

39 Gary, R. E., Jr. & Foster, W. A. Diel timing and frequency of sugar feeding in the mosquito *Anopheles gambiae*, depending on sex, gonotrophic state and resource availability. Med. Vet. Entomol. 20, 308–316, doi:10.1111/j.1365-2915.2006.00638.x (2006).

40 Jones, M. D. R., Hill, M. & Hope, A. M. The circadian flight activity of the mosquito *Anopheles gambiae*: phase setting by the light regime. J. Exp. Biol. 47, 503–511 (1967).

41 Lima-Camara, T. N., Lima, J. B., Bruno, R. V. & Peixoto, A. A. Effects of insemination and blood-feeding on locomotor activity of *Aedes albopictus* and *Aedes aegypti* (Diptera: Culicidae) females under laboratory conditions. Parasit. Vectors 7, 304–312, doi:10.1186/1756-3305-7-304 (2014).

42 Yee, W. L. & Foster, W. A. Diel sugar-feeding and host-seeking rhythms in mosquitoes (Diptera: Culicidae) under laboratory conditions. J. Med. Entomol. 29, 784–791, doi:10.1093/jmedent/29.5.784 (1992).

43 Jones, M. D. R. The programming of circadian flight-activity in relation to mating and the gonotrophic cycle in the mosquito, *Aedes aegypti*. Physiol. Entomol. 6, 307–313 (1981).

44 Taylor, B. & Jones, M. D. The circadian rhythm of flight activity in the mosquito *Aedes aegypti* (L.). The phase-setting effects of light-on and light-off. J. Exp. Biol. 51, 59–70 (1969).

45 Lees, R. S. et al. Review: Improving our knowledge of male mosquito biology in relation to genetic control programmes. Acta Trop. 132 Suppl, S2–11, doi: 10.1016/j.actatropica.2013.11.005 (2014).

46 Oliva, C. F., Damiens, D., Vreysen, M. J., Lemperiere, G. & Gilles, J. Reproductive strategies of *Aedes albopictus* (Diptera: Culicidae) and implications for the sterile insect technique. PLoS One 8, e78884, doi:10.1371/journal.pone.0078884 (2013).

47 Lima-Camara, T. N. Activity patterns of *Aedes aegypti* and *Aedes albopictus* (Diptera: Culicidae) under natural and artificial conditions. Oecol. Aust. 14, 737–744 (2010).

48 Gerberg, E. J., Barnard, D. R. & Ward, R. A. Manual for Mosquito Rearing and Experimental Techniques. (American Mosquito Control Association, Inc., 1994).

49 Donelson, N.C. et al. Correction: high-resolution positional tracking for long-term analysis of *Drosophila* sleep and locomotion using the “Tracker” program. PloS One 7, e37250, doi.org/10.1371/journal.pone (2012).

50 Levine, J. D., Funes, P., Dowse, H. B., Hall, J. C. Signal analysis of behavioral and molecular cycles. BMC Neurosci. 3, 1–25 (2002).

